# The highly conserved stem-loop II motif is dispensable for SARS-CoV-2

**DOI:** 10.1101/2023.03.15.532878

**Authors:** Hongbing Jiang, Astha Joshi, Tianyu Gan, Andrew B Janowski, Chika Fujii, Traci L Bricker, Tamarand L Darling, Houda H. Harastani, Kuljeet Seehra, Hongwei Chen, Stephen Tahan, Ana Jung, Binita Febles, Joshua A Blatter, Scott A Handley, Bijal A Parikh, David Wang, Adrianus CM Boon

## Abstract

The stem-loop II motif (s2m) is a RNA structural element that is found in the 3’ untranslated region (UTR) of many RNA viruses including severe acute respiratory syndrome coronavirus 2 (SARS-CoV-2). Though the motif was discovered over twenty-five years ago, its functional significance is unknown. In order to understand the importance of s2m, we created viruses with deletions or mutations of the s2m by reverse genetics and also evaluated a clinical isolate harboring a unique s2m deletion. Deletion or mutation of the s2m had no effect on growth *in vitro*, or growth and viral fitness in Syrian hamsters *in vivo*. We also compared the secondary structure of the 3’ UTR of wild type and s2m deletion viruses using SHAPE-MaP and DMS-MaPseq. These experiments demonstrate that the s2m forms an independent structure and that its deletion does not alter the overall remaining 3’UTR RNA structure. Together, these findings suggest that s2m is dispensable for SARS-CoV-2.

**IMPORTANCE:** RNA viruses, including severe acute respiratory syndrome coronavirus 2 (SARS-CoV-2) contain functional structures to support virus replication, translation and evasion of the host antiviral immune response. The 3’ untranslated region of early isolates of SARS-CoV-2 contained a stem-loop II motif (s2m), which is a RNA structural element that is found in many RNA viruses. This motif was discovered over twenty-five years ago, but its functional significance is unknown. We created SARS-CoV-2 with deletions or mutations of the s2m and determined the effect of these changes on viral growth in tissue culture and in rodent models of infection. Deletion or mutation of the s2m element had no effect on growth *in vitro*, or growth and viral fitness in Syrian hamsters *in vivo*. We also observed no impact of the deletion on other known RNA structures in the same region of the genome. These experiments demonstrate that the s2m is dispensable for SARS-CoV-2.

## BACKGROUND

The RNA genome of severe acute respiratory syndrome coronavirus 2 (SARS-CoV-2) is approximately 30,000 nucleotides in length^1^. It consists of a 5’ untranslated region (UTR), coding sequences for structural and non-structural proteins, and a 3’ UTR. The 3’ UTR contains highly structured RNA elements such as stem-loop sequence (SL1), bulged stem-loop (BSL), pseudoknot (PK), and a hypervariable region (HVR) which have been suggested to function in viral genome replication, transcription, and viral protein translation^2,3^. SARS-CoV-2, SARS-CoV-1, and other members of *Sarbecovirus* lineage in the *Betacoronavirus* genus, as well as some members of the *Gammacoronavirus* and *Deltacoronavirus* genus encode a stem-loop II motif (s2m) within the terminal portion of HVR in the 3’ UTR^4,5^ (**Fig. 1A**). In contrast, seasonal human coronaviruses (HKU1, 229E, OC43, and NL63) and middle eastern respiratory syndrome coronavirus (MERS-CoV) do not contain an s2m in their genomes^4^. The s2m element has also been detected in members of the *Astroviridae, Caliciviridae, Coronaviridae, Picornaviridae*, and *Reoviridae* viral families, all with highly conserved nucleotide sequences of 39-43 nucleotides in length^4,6–9^. Currently, the function of the s2m for the viral lifecycle is poorly understood. Phylogenetic distribution suggests horizontal acquisition of the s2m at different timepoints, and maintenance of the element suggests that it may confer a fitness advantage^4,6^. The X-ray crystal structure of the s2m element from SARS-CoV-1 demonstrates a stem-loop secondary structure and a tertiary structure that includes a 90° kink in the helix axis resulting in additional tertiary interactions^7^. The secondary structure determination by NMR and probing methods for the SARS-CoV-2 s2m element revealed two stem structures separated by an internal asymmetric loop^10–15^. Antisense oligonucleotides targeting the s2m reduced viral replication for SARS-CoV-2 and classic human astrovirus 1 (HAstV1) replicons^16^. The SARS-CoV-2 s2m was shown to dimerize and interact with host cellular microRNA 1307-3p^17^. These results suggest that the secondary structure of the s2m is conserved and potentially important for viral replication or other host-virus interactions.

**Figure 1:**
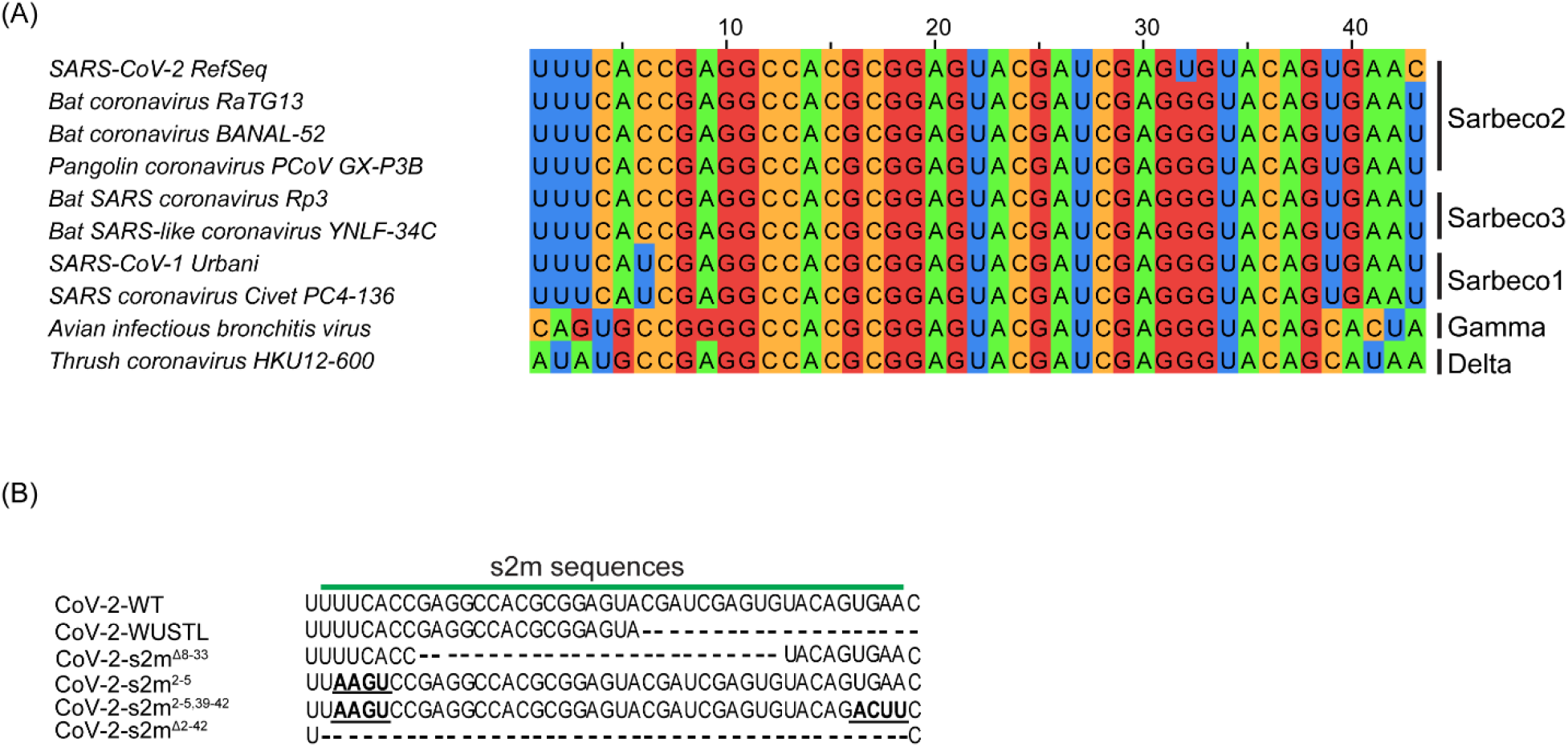
s2m nucleotide sequence alignment. (A) Multiple sequence alignment of representative Coronaviruses in the *Betacoronavirus*, *Gammacoronavirus*, and *Deltacoronavirus* genus that encode an s2m. (B) Sequence alignment of the s2m element found in SARS-CoV-2 isolates. SARS-CoV-2-WT is the original SARS-CoV-2 isolate (NC_045512.2), SARS-CoV-2-WUSTL is the virus isolated from the patient in the St Louis Area (2019-nCoV/WUSTL_000226/2020), SARS-CoV-2-s2m^Δ8-33^ is the circulating s2m deletion mutant, SARS-CoV-2-s2m^2-5^ is the engineered s2m stem structure mutant, SARS-CoV-2-s2m^2-5,39-42^ is the engineered s2m stem structure mutation revertant, and SARS-CoV-2-s2m^Δ2-42^ is the engineered s2m deletion mutant.

Interestingly, the SARS-CoV-2 genomes encode a uracil residue at position 32 in the s2m (position 29,758 in the virus genome) that is distinct from all known s2m sequences in other viruses and is predicted to perturb the secondary structure^10,11,18–20^. Additional genetic variants or deletions of the s2m have also been periodically detected in clinical isolates prior to the emergence of the BA.2 Omicron lineage of SARS-CoV-2^20–23^. However, SARS-CoV-2 variants that emerged after December 2021 contain a 26-nucleotide deletion of the s2m element (**Fig S1**)^24^. Combined, these data suggests that the s2m element has minimal or no impact on the lifecycle of SARS-CoV-2.

Here, we determined the functional significance of s2m *in vitro* and *in vivo* using recombinant SARS-CoV-2 viruses or natural isolates with mutations or deletions in the s2m element in the 3’ UTR of the genome. We also determined the 3’ UTR RNA structure of SARS-CoV-2 in the presence or absence of the s2m element. We show that deletion of s2m in SARS-CoV-2 has no impact on the viral fitness or 3’ UTR structure of SARS-CoV-2.

## RESULTS

### Natural varation and deletion of s2m found in SARS-CoV-2 circulating strains

The original SARS-CoV-2 virus genome (reference genome: NC 045512) contains an s2m element in the 3’ UTR region, which is similar to other Sarbecoviruses in the *Betacoronavirus* genus, and some members of the *Gammacoronavirus* and *Deltacoronavirus* genus (**Fig. 1A**). Interestingly, the SARS-CoV-2 s2m encodes a uracil at position 32 of the s2m (position 29758 in the reference genome) while essentially all other s2m sequences in coronaviruses known to date contain guanine (**Fig. 1A**). In the s2m element of SARS-CoV-1, this position forms a G-C base pair as determined by X-ray crystal structure^7^ and by RNAfold prediction. To further examine the variation at this position, we analyzed 1,705,180 complete SARS-CoV-2 genomes uploaded to the NCBI database from January 2020 to December 2022. Of those sequences that contained a complete s2m element, only eight genomes contained guanine at this position. An additional 18 had a cytosine residue and one genome contained an adenine at position 32. We also noticed in the multiple sequence alignment that position 9 can be variable between viruses within the coronavirus family (**Fig. 1A**). SARS-CoV-1 and SARS-CoV-2 encode an adenine while avian infectious bronchitis virus encodes a guanine. This position is not predicted to form any base pair, but has been identified to form potential long-distance tertiary interactions with nucleotide 30 in the SARS-CoV-1 s2m crystal structure^7^.

Although the s2m element is relatively conserved in the genome of SARS-CoV-2 between the start of the pandemic and early 2022, several genomes were found to have a partial or complete deletion of the s2m element^23^. Our analysis of the 1,705,180 complete SARS-CoV-2 genomes also revealed the emergence of SARS-CoV-2 lineages with a 26-nucleotide deletion (position 8 to 33) in the s2m element (**Fig. 1B and S1A**). The genomes that contain this deletion mainly belong to BA.2 lineage (Pango Lineages) of SARS-CoV-2 which includes BA.2.75, BA.4, BA.5, and the recent BQ.1 and XBB.1 variants of SARS-CoV-2 (**Fig. S1B**).

### The s2m element is dispensable for SARS-CoV-2 *in vitro*

To determine the importance of the s2m in SARS-CoV-2 virus lifecycle, we recombinantly generated in the reference backbone, wild type (CoV-2-s2m-WT) and three mutant viruses with mutations or deletions in the s2m region using the reverse genetic system described in **Fig. S2**. To remove the s2m element in the 3’ UTR of SARS-CoV-2, we deleted nucleotides 2-42 in the s2m region (CoV-2-s2m^Δ2-42^, **Fig. 1B**). We also created a mutant that contained four consecutive nucleotide substitutions in the stem region of the s2m (CoV-2-s2m^2-5^) that is predicted to disrupt the secondary structure as well as a revertant mutant (CoV-2-s2m^2-5, 39-42^) that contained four additional compensatory substitutions that restored the predicted stem region and the SARS-CoV-2 s2m secondary structure (**Fig. 1B**). All mutant SARS-CoV-2 genomes yielded infectious viral particles that could be propagated in Vero-hTMPRSS2 cells. We confirmed the presence of the engineered mutations and the absence of spontaneous mutations in the SARS-CoV-2 s2m by sequencing. We next tested whether there were any defects in the growth rate of the mutants by conducting multi-step growth curves (**Fig. 2A**). There was no significant difference between CoV-2-s2m-WT and mutant viruses at any timepoint in Vero-hTMPRSS2 (African green monkey) cells (two-way ANOVA F[3,20] = 1.02, *P* = 0.4) or Calu-3 (human) cells (two-way ANOVA F[3,20] = 0.48, *P* = 0.7), suggesting that the s2m element is not required for SARS-CoV-2 lifecycle *in vitro* (**Fig. 2B**).

**Figure 2:**
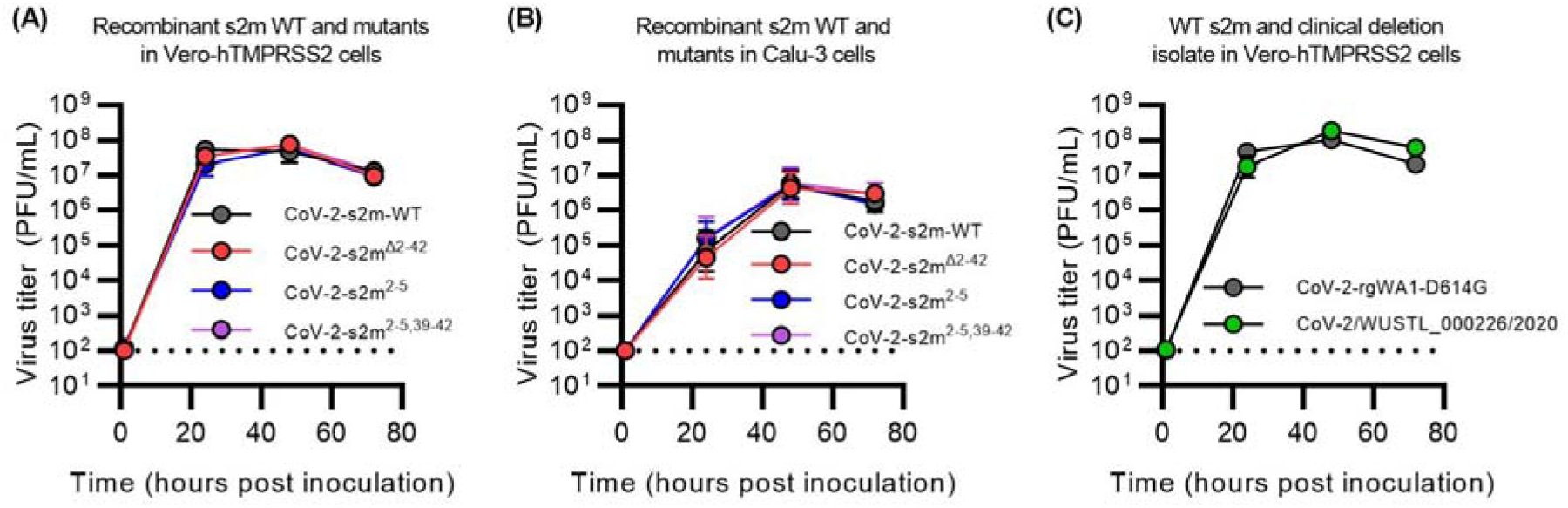
The s2m is dispensable for SARS-CoV-2 *in vitro*. (**A-B**) Multi-step growth curve of CoV-2-s2m-WT and mutants in (**A**) Vero-hTMPRSS2 and (**B**) Calu-3 cells. Mutant strains include CoV-2-s2m^Δ2-42^ which contains a deletion of the s2m element, CoV-2-s2m^2-5^ which contains four consecutive substitutions of the stem of the s2m, and CoV-2-s2m^2-5,39-42^ which contains complementary substitutions predicted to restore the secondary structure of the s2m. Infectious virus titer measured in plaque forming units per mL (PFU/mL) at 0, 24, 48, and 72 hours post-inoculation. No difference in the viral titer was detected by a two-way ANOVA with post-hoc testing by Dunnett’s multiple comparison test between CoV-2-s2m-WT and all mutants for Vero-hTMPRSS2 F(3,20)= 1.02, P= 0.40, and for Calu-3 cells F(3,20)= 0.48, P= 0.7). (**C**) Multi-step growth curve of the infectious viral titer using a clinical isolate of SARS-CoV-2 containing a partial deletion of the s2m element (WUSTL_000226/2020), compared to a WA1-strain of SARS-CoV-2 with a D614G mutation. Viral titers were measured at 0, 24, 48, and 72 hours post-inoculation and no difference in titers was detected by a two-way ANOVA with post-hoc testing by Sidak’s multiple comparison test (F(1,20)= 2.34, P= 0.16). For all graphs, geometric means ± geometric standard deviations are depicted. Dotted line is the limit of detection of the assay.

### Growth of a clinical SARS-CoV-2 isolate with a deletion in the s2m

During genomic surveillance for SARS-CoV-2 variants in the St. Louis area, USA, we identified one genome (SARS-CoV-2/WUSTL_000226/2020) that contained a deletion of 27 nucleotides that removes positions 22-43 of the s2m and additional nucleotides in the 3’ UTR **(Fig. 1B**). This mutant belongs to the B1.2 lineage (Pango Lineages)^25^ which contains the D614G variant in the spike protein, and has 99.8% nucleotide identity compared to the reference SARS-CoV-2 genome (NC_045512.2). We were able to culture WUSTL_000226/2020 and verified that the recovered virus maintained the deletion by sequencing. In a multi-step growth curve, we did not observe any significant difference in virus titer between WUSTL_000226/2020 and a recombinant WA1-strain of SARS-CoV-2 engineered with D614G mutation at any timepoint (two-way ANOVA F[1,10] = 2.3, *P* = 0.16; **Fig. 2C**).

### The s2m element is dispensable for SARS-CoV-2 infection and replication *in vivo*

We next determined if the SARS-CoV-2 s2m was important *in vivo* using the Syrian hamster model. Hamsters were intranasally inoculated with 1,000 plaque forming units of the CoV-2-s2m-WT and mutant viruses and weights were recorded daily for six days. Nasal washes and lungs were collected 3 and 6 days post infection and infectious virus titer and viral RNA load was quantified by plaque assay and RT-qPCR respectively. No difference in weight loss was observed between the hamsters inoculated with the CoV-2-s2m-WT and deletion or mutant s2m-containing viruses (mixed-effect model with Geisser-Greenhouse correction F[3,44] = 1.26, *P* = 0.30). (**Fig. 3A**). No significant differences in infectious virus (lung tissue Kruskal-Wallis H(3) = 0.62, *P* = 0.89), viral RNA (lung tissue Kruskal-Wallis H(3) = 2.0, *P* = 0.57; nasal wash Kruskal-Wallis H(3) = 2.2, *P* = 0.54) were detected for CoV-2-s2m^Δ2-42^, CoV-2-s2m^2-5^, and CoV-2-s2m^2-5,39-42^ compared to CoV-2-s2m-WT (**Fig. 3B-D**). At six days post-inoculation, no difference in infectious virus was detected in the lungs of the mutant SARS-CoV-2 infected hamsters compared to CoV-2-s2m-WT (Kruskal-Wallis H(3) = 5.5, *P* = 0.14; **Fig. 3E**). The viral RNA load with CoV-2-s2m^Δ2-42^, CoV-2-s2m^2-5^, and CoV-2-s2m^2-5,39-42^ also showed no difference compared to CoV-2-s2m-WT from the lungs (Kruskal-Wallis H(3) = 6.2, *P* = 0.10) and nasal washes (Kruskal-Wallis H(3) = 2.8, *P* = 0.42; **Fig. 3F-G**). Combined, these data suggest that the original SARS-CoV-2 virus does not require the s2m for growth *in vitro* or *in vivo*.

**Figure 3:**
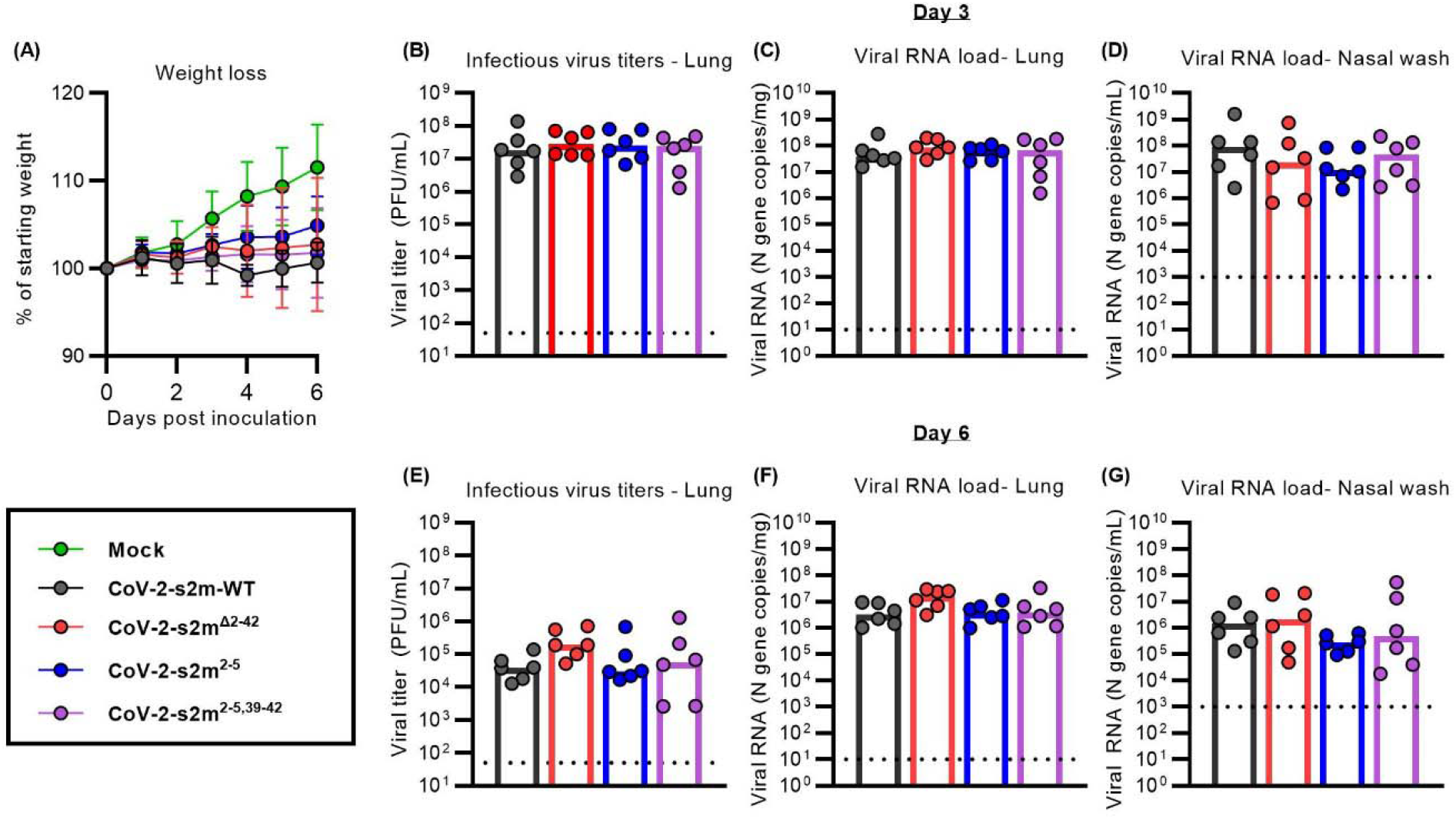
The s2m is dispensable for SARS-CoV-2 *in vivo*. Intranasal inoculation of hamsters with CoV-2-s2m-WT and mutants including CoV-2-s2m^Δ2-42^ which contains a deletion of the s2m, CoV-2-s2m^2-5^ that contains four mutations of the stem of the s2m, and CoV-2-s2m^2-5,39-42^ that contains complementary mutations predicted to restore the secondary structure of the s2m. Hamsters were sacrificed at days 3 and 6. (**A**) Mean hamster weight as percent of starting weight is graphed with error bars representing standard deviations. The weight of the hamsters infected with CoV-2-s2m-WT and mutant SARS-CoV-2 viruses were compared and no difference was detected using a mixed effect model with Geisser-Greenhouse correction (F[3,44] =1.26, P= 0.30). There was no difference in (**B**) Lung infectious viral titer (Kruskal-Wallis H(3)= 0.62 P= 0.89), (**C**) lung viral RNA (Kruskal-Wallis H(3)= 2.0 P= 0.57), and (**D**) nasal wash viral RNA (Kruskal-Wallis H(3)= 2.2 P= 0.54) from day 3 were identified comparing CoV-2-s2m-WT and s2m mutant viruses. At 6 days post inoculation, there was also no differences in (**E**) lung infectious viral titer (Kruskal-Wallis H(3)= 5.5 P= 0.14), (**F**) lung viral RNA (Kruskal-Wallis H(3)= 6.2 P= 0.10), and **(G)** nasal wash viral RNA (Kruskal-Wallis H(3)= 2.8 P= 0.42) comparing CoV-2-s2m-WT to mutant s2m viruses. For all graphs B-G, each dot is one animal. Dotted line represents the limit of detection of the respective assay.

### The s2m element is dispensable for SARS-CoV-2 viral fitness *in vivo*

To further determine if the SARS-CoV-2 s2m element has any effect on viral fitness *in vivo*, we designed a viral competition assay using the CoV-2-s2m-WT and the CoV-2-s2m^Δ2-42^ virus in Syrian hamsters. CoV-2-s2m-WT and the CoV-2-s2m^Δ2-42^ mutant virus were mixed at a 1:1 ratio and inoculated intranasally into Syrian hamsters. The ratio CoV-2-s2m-WT to CoV-2-s2m^Δ2-42^ mutant virus in the inoculum was determined by RT-PCR on the 3’ UTR following RNA extraction of RNase treated virus (**Fig. 4A**). Three days post infection (dpi), the lungs and nasal washes were collected and the genome copy number of CoV-2-s2m-WT to CoV-2-s2m^Δ2-42^ was measured by RT-PCR on RNA extracted from these tissues and samples. The relative replicative fitness of CoV-2-s2m^Δ2-42^ to CoV-2-s2m-WT was calculated for each sample. The mean relative replicative fitness of CoV-2-s2m^Δ2-42^ to CoV-2-s2m-WT was 1.21 and 1.10 in the lung and nasal washes respectively (**Fig. 4B**). This ratio was similar to that of the input, indicating that the CoV-2-s2m-WT virus has no fitness advantage over the CoV-2-s2m^Δ2-42^ mutant virus *in vivo* in Syrian hamsters.

**Figure 4:**
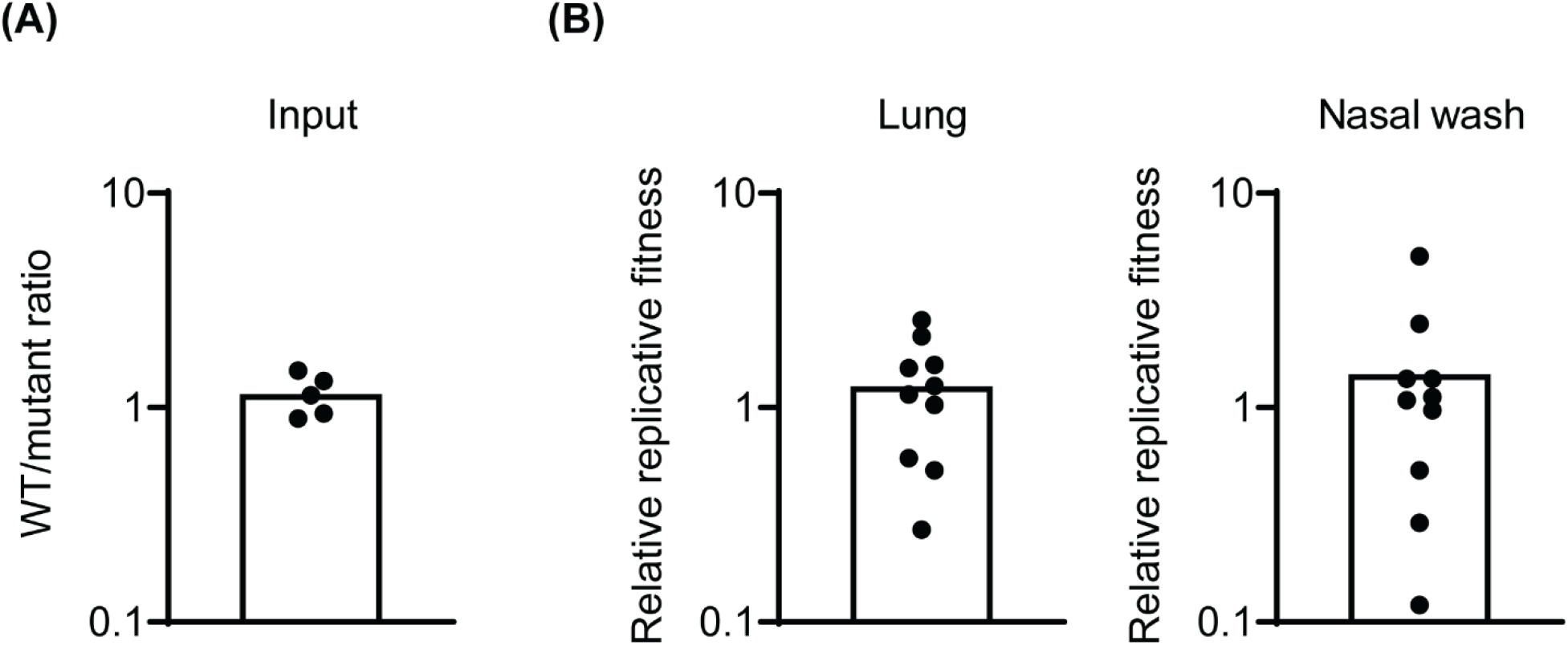
The s2m is dispensable for SARS-CoV-2 viral fitness in Syrian hamster. The fitness of the CoV-2-s2m^Δ2-42^ virus was assessed *in vivo* in Syrian hamsters. Hamsters were inoculation intranasally with 1:1 mixture of CoV-2-s2m-WT and CoV-2-s2m^Δ2-42^ virus and lungs and nasal washes were collected 3 days post infection. The genome copy of CoV-2-s2m-WT and CoV-2-s2m^Δ2-42^ in the inoculum and infected tissues were measured by RT-PCR. The ratio of CoV-2-s2m-WT to CoV-2-s2m^Δ2-42^ in the inoculum was calculated (**A**) and the replicative fitness of the CoV-2-s2m^Δ2-42^ to CoV-2-s2m-WT virus in lungs and nasal washes were calculated (**B**). Individual values were shown as dots and mean values were plotted by box.

### Structural analysis of the 3’ UTR of SARS-CoV-2

In order to determine the RNA secondary structure of the 3’ UTR in the SARS-CoV-2 genome and the impact of the s2m deletion on the secondary structure, we performed SHAPE-MaP and DMS-MaPseq RNA structure probing studies on purified CoV-2-s2m-WT and CoV-2-s2m^Δ2-42^ virus using NAI and DMS respectively.

RNA structure predictions, using the SHAPE-MaP reactivity data as constraints, identified the bulged-stem loop (BSL), stem loop 1 (SL1), pseudoknot, and a bulge stem that includes the hypervariable region (HVR), the s2m element, and the octanucleotide motif (ONM) in the 3’ UTR of CoV-2-s2m-WT (**Fig. 5A**). Interestingly, DMS-MaPseq predicted a similar secondary structure with the BSL, SL1, HVR, s2m element, and the ONM motif (**Fig. 5B**). Overall, the SHAPE-MaP and DMS-MaPseq reactvities were in good agreement with the predicted structure (**Fig. S3**) and with previous findings^11–15^ Structure prediction using our SHAPE-MaP data also predicted the formation of a pseudoknot between the base of the BSL and the loop of SL1 (**Fig. 5A**). This pseudoknot was not predicted using DMS-MaPseq reactivity data (**Fig. 5B**). Using Superfold, we found that the 3’ UTR is highly structured and all the known structured elements have high base-pairing probablities shown by green color (**Fig. S3**).

**Figure 5:**
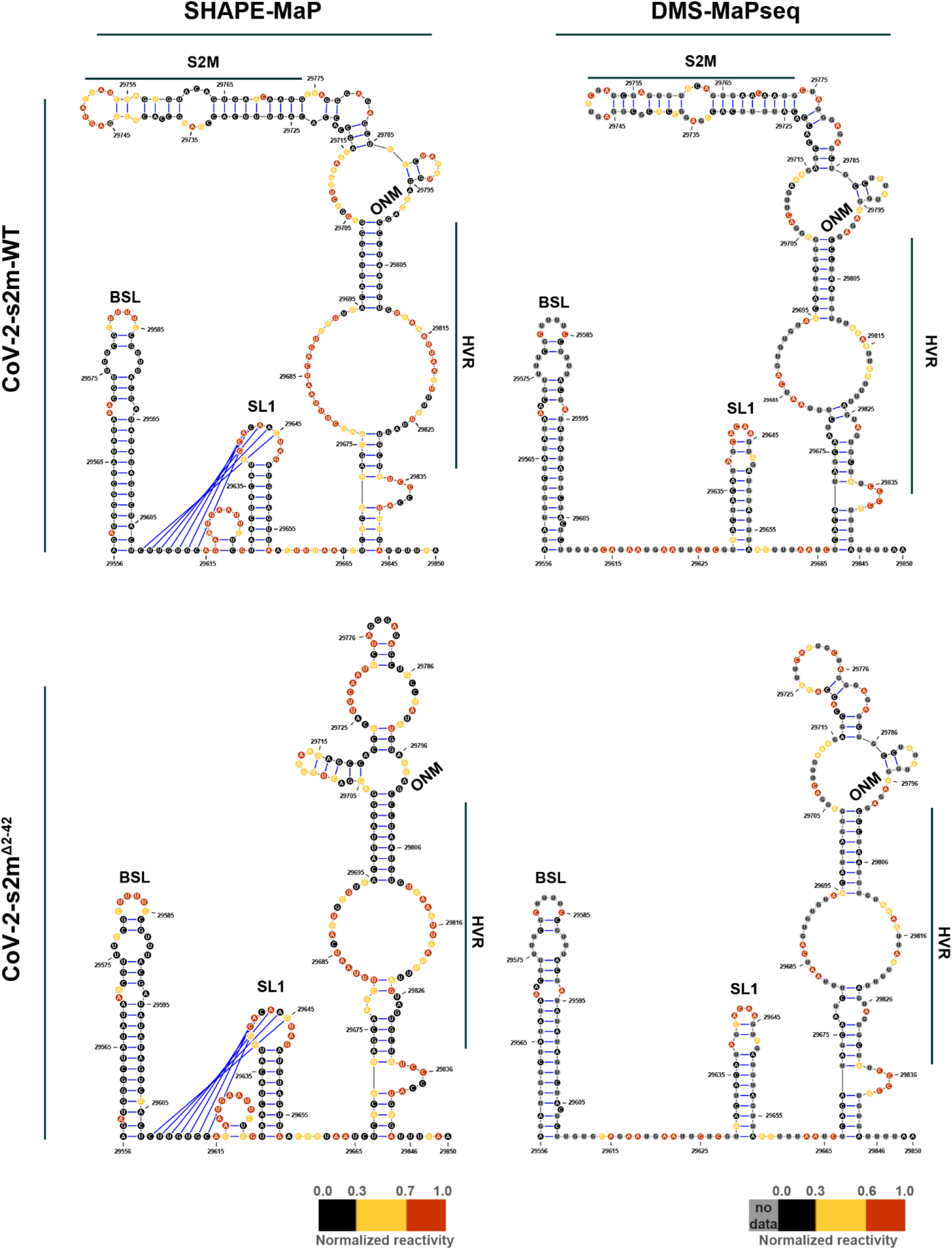
Structure prediction for 3’UTR suggests a presence of conserved structural elements. Predicted RNA secondary structure for CoV-2-s2m-WT using SHAPE-MaP (**A**) and DMS-MaPseq (**B**) reactivity profiles. Predicted RNA secondary structure for CoV-2-s2m^Δ2-42^ using SHAPE-MaP (**C**) and DMS-MaPseq (**D**) reactivity profiles. Each nucleotide is colored by their normalized reactivity. BSL, bulged-stem loop; SL1, stem loop 1; HVR, hypervariable region; ONM, the octanucleotide motif. Blue lines indicate the pseudoknot region.

SHAPE-MaP on the 3’ UTR of CoV-2-s2m^Δ2-42^ revealed a very similar reactivity profile and the predicted structure contained the BSL, SL1, HVR, and ONM region. Comparison between the CoV-2-s2m-WT and CoV-2-s2m^Δ2-42^ reactivities showed a high correlation (R^2^ = 0.88). Similar to CoV-2-s2m-WT, the SL1 region of CoV-2-s2m^Δ2-42^ was also predicted to form a pseudoknot with the base of the BSL region. Analogous to the SHAPE-MaP data, the DMS-MaPseq reactivity and predicted structure between CoV-2-s2m-WT and CoV-2-s2m^Δ2-42^ showed a high correlation (R^2^ = 0.92). The only difference in reactivity and structure was observed near the s2m region that was deleted in the CoV-2-s2m^Δ2-42^ virus. These data suggest that 3’ UTR of SARS-CoV-2 is a rigid structure and that deletion of the s2m region did not change the overall 3’ UTR structure.

## DISCUSSION

Although the crystal structure of the conserved s2m RNA element was solved for SARS-CoV-1 in 2005^7^ and many recent studies suggested an important role of the s2m structure for SARS-CoV-2, no functional genetic studies on the s2m element in the coronavirus lifecycle have been performed^4,6–9,26^. Here, we demonstrated that the deletion or mutation of the s2m element in the original strain of SARS-CoV-2 did not impact growth *in vitro* or viral fitness *in vivo*, and had minimal effect on the predicted RNA secondary structure of the 3’ UTR of SARS-CoV-2. These results suggest that the s2m structure is not essential for SARS-CoV-2.

Based on the presence and conservation of the s2m element in different viral families, it has been hypothesized to be beneficial for the virus. However, our results suggest that the function of the s2m element is dispensable in the SARS-CoV-2 genome, perhaps due to redundancy with another uncharacterized virus-derived element, whether a viral protein or RNA motif. Investigating the signifance of the s2m element in related Sarbecoviruses, including SARS-CoV-1, RaTG13 and pangolin coronaviruses, is needed to better understand its role in coronavirus biology. Given the genetic diversity within the coronavirus family, it is unclear if the s2m is important or redundant in other coronaviruses. SARS-CoV-2 is the only coronavirus that contains a uracil at position 32, while all others have a guanine at this site (**Fig. 1A**)^27^. The presence of this uracil may represent a unique and recent evolutionary event for the s2m that is specific to SARS-CoV-2 lineage, and therefore the biology of the SARS-CoV-2 s2m may not apply to other coronaviruses. Complete genome analysis of natural isolates of SARS-CoV-2 also found that a small subset contained mutations or deletions in the s2m region^20–23^. One of these isolates was found by our group and we showed no difference in growth potential in two different cell lines compared to a related SARS-CoV-2 with the s2m element intact. While it is possible that these rare deletions were detected because of the unprecedented amount of genome sequencing that was done during the pandemic, it is also possible that this was an early indication that this region was under neutral or negative selective pressure facilitating the emergence of the Omicron lineage with a 26-nucleotide deletion in early 2022 (**Fig S1A-B**). Analogous to SARS-CoV-2, the genomes of several seasonal coronaviruses and MERS also do not contain a s2m element^4^. Whether this is an adaptation of coronaviruses to the human host remains to be investigated.

Our RNA structure modeling of the 3’ UTR present in virions indicated the presence of several conserved structural elements including the BSL, SL1, and a bulge stem that includes the HVR, the s2m element, and the ONM. Overall, our model is similar to those identified by others^11–15^. The BSL, SL1, and s2m form a stem loop structure. The bulge in the BSL motif was found to have low SHAPE-MaP reactivity which can be explained by the binding of a viral or host protein as suggested previously^11^. The predicted structure of the HVR was different between the two probing methods and between different studies. It is mostly a single stranded region with high reactivity bases between two structured regions, which may explain why it tolerates the presence of multiple mutations and is not essential for viral RNA synthesis^21,28^. ONM (5’-GGAAGAGC-3’) is known to be a single-stranded region with a critical biological function^28^. In our model, the first two nucleotides (GG) of the ONM form the base of a small hairpin. Currently, it is not known if this hairpin is an artifact of the RNA folding software or whether it is present in the 3’ UTR of SARS-CoV-2. Interestingly, the reactivity of the ONM region was overall lower in SHAPE-MaP compared to DMS-MaPseq, which can be explained by the binding of a viral or host protein to the ONM. The predicted structure of the s2m element was similar between SHAPE-MaP and DMS-MaPseq. Compared to a previously determined crystal structure of s2m in SARS-CoV-1^7^, the main differences are found near the top of the s2m element, with a predicted extended loop in the s2m of SARS-CoV-2. This is potentially caused by the uracil at position 32 resulting in different base pairings. This uracil was found to be moderately reactive by SHAPE-MaP and therefore predicted to interact weakly with the adenosine at position 14 of the s2m element. The weak interaction is supported by DMS-MaPseq which showed a moderate reactivity of adenosine at position 14 (DMS does not modify uracil residues).

In contrast to DMS-MaPseq, SHAPE-MaP predicted the formation of a tertiary pseudoknot structure between the base of the BSL and the loop of SL1. The confidence of this tertiary structure prediction is relatively low since many of the involved nucleotides are highly reactive by SHAPE-MaP or DMS-MaPseq. As such, it is possible that no pseudoknot is formed as suggested by others^11^. Alternatively, this region of the 3’ UTR switches between a tertiary structure (pseudoknot) and a free SL1 structure in SARS-CoV-2. This model is supported by recent studies on Murine Hepatitis virus that suggests that both structures contribute to viral replication and may function as molecular switches in different steps of RNA synthesis^2^.

The comparison between the predicted RNA structures of the CoV-2-s2m-WT and CoV-2-s2m^Δ2-42^ 3’ UTR demonstrated nearly identical structures. Only the region containing the s2m element was significantly different between the two viruses. These data suggests that the s2m hairpin structure does not interact with any other elements in the 3’UTR region of SARS-CoV-2 as predicted previously by computational analysis^29^. Taken together, our RNA structural analysis suggests that there is no impact on the overall 3’ UTR RNA structure upon deletion of the s2m element, as found in the majority of the SARS-CoV-2 virus isolated since March 2022. However, the lack of interactions between s2m and the rest of the 3’ UTR we observed in SARS-COV-2 by SHAPE-MaP and DMS-MaPseq may differ considerably between different viruses and virus families. The presence and conservation of s2m in different viral families suggest that it may still habor important functions for virus lifecycle in viruses other than SARS-CoV-2 and warrants further investigation.

### Limitations of the study

We note several limitations of our study. (a) The RNA structure analyses were done on purified virus and not on viral genomes inside immunocompetent cells. While our structure predictions of the 3’ UTR are similar to those obtained from infected cells, it is possible that the SHAPE-MaP and DMS-MaPseq reactivity is different in primary human nasal or bronchial epithelial cells. (b) SHAPE-MaP and DMS-MaPseq are low resolution probing methods that average out the reactivity of particular nucleotides, potentially obscuring alternative or higher order structures. (c) The transmission potential of CoV-2-s2m-WT and s2m mutant viruses was not assessed. Airborne transmission of the CoV-2-s2m-WT viruses is inefficient and while we did detect airborne transmission of both CoV-2-s2m-WT and mutant virus, the results were inconclusive and therefore not included here. (d) It is possible that the significance of the s2m element is cell-line or host dependent. While we did not test primary differentiated airways epithelial cell cultures or additional animal models (mouse or non-human primates), the emergence and dominance of SARS-CoV-2 viruses lacking the s2m element supports our conclusion that s2m is dispensible for the SARS-CoV-2 life cycle. (e) The impact of the s2m deletion and mutations on lung pathology was not fully assessed. We opted to inoculate the animals with a relatively low dose to allow the detection of replication differences. However, these lower doses do not induce large amounts of weight loss or lung pathology. Future studies could detect transcriptional differences in the lungs of hamsters infected with CoV-2-s2m-WT or s2m deletion viruses in order to identify a role of s2m in modulating host responses.

Overall, we have found that the s2m element is not critical for SARS-CoV-2 virus lifecycle *in vitro* or viral fitness *in vivo*. Further studies are needed to define the mechanistic basis as to why a highly conserved RNA element has no critical functional roles in the viral lifecycle.

## MATERIALS AND METHODS

### Cell culture conditions

BHK-21cells were maintained in Dulbecco’s Modified Eagle Medium (DMEM) with L-glutamine (Gibco) supplemented with 10% fetal bovine serum (FBS) and 100 units/mL penicillin/streptomycin and incubated at 37°C and 5% CO_2_. Vero cells expressing human TMPRSS2 (Vero-hTMPRSS2)^30^ or human ACE2 and human TMPRSS2 (Vero-hACE2-hTMPRSS2, gift from Drs. Graham and Creanga at NIH) and BSR cells (a clone of BHK-21) were cultured and maintained in DMEM supplemented with 5% FBS and 100 units/mL of penicillin and streptomycin. Vero-hTMPRSS2 and Vero-hACE2-hTMPRSS2 cells were maintained by selection with 5 μg/mL Blasticidin or 10 μg/mL puromycin respectively.

### SARS-CoV-2 reverse genetics system

All work with potentially infectious SARS-CoV-2 particles was conducted under enhanced biosafety level 3 (BSL-3) conditions and approved by the institutional biosafety committee of Washington University in St. Louis. The prototypic SARS-CoV-2 genome (reference genome NC_045512.2) was split into 7 fragments (**Fig S1**), named A to G, and each DNA fragment was commercially synthesized (GenScript). A T7 promoter sequence was introduced at the 5’ end of fragment A, and a poly-A sequence of 22 adenosines was introduced at the 3’ end of fragment G. In addition, NotI and SpeI sites were introduced at the 5’ end of fragment A before the T7 promoter and the 3’ end of fragment G after the poly-A sequence, respectively. To ensure seamless assembly of the full virus DNA genome, the 3’ end of fragment A, both ends of fragments B-F and the 5’ end of fragment G were appended by class II restriction enzyme recognition sites (BsmBI and BsaI) (**Fig S1**). Fragments A and C-G were cloned into plasmid pUC57 vector and amplified in *E. Coli* DH5α strain. The bacteria toxic fragment B was cloned into low copy inducible BAC vector pCCI and amplified through plasmid induction in EPI300. Low copy plasmids were extracted by NucleoBond Xtra Midi kit (MACHEREY-NAGEL) with the other plasmids were extracted by plasmids midi kit (QIAGEN) according to the manufacturers’ protocols. To generate mutant s2m sequences, the SARS-CoV-2 s2m sequence in the G fragment was mutated by site-directed mutagenesis (**Fig. 1B**). Mutations and deletions in the s2m element were confirmed by Sanger sequencing. To assemble the full-length SARS-CoV-2 genome, each DNA fragment in plasmid was digested by corresponding restriction enzymes, DNA fragments were recovered by gel purification columns (New England Biolabs), and the seven fragments were ligated at equal molar ratios with 10,000 units of T4 ligase (New England Biolabs) in 100 μL at 16°C overnight. The final ligation product was incubated with proteinase K in the presence of 10% SDS for 30 minutes, extracted twice with equal volume of phenol:chloroform:isoamyl alcohol (25:24:1; ThermoFisher), isopropanol precipitated, air dried and resuspended in RNase/DNase free water. The DNA was analyzed on a 0.6% agarose gel. Full length genomic SARS-CoV-2 RNA was *in vitro* transcribed using mMESSAGE mMACHINE T7 Ultra Transcription kit (Invitrogen) following the manufacturer’s protocol. Four μg of DNA template was added to the reaction mixture, supplemented with GTP (7.5 μL per 50 μL reaction). *In vitro* transcription was done overnight at 32°C. Afterwards, the template DNA was removed by digestion with Turbo DNase for 30 minutes at 37°C. The *in vitro* transcript (IVT RNA) mixture was used directly for electroporation. To further enhance rescue of recombinant virus, we also generated SARS-CoV-2 nucleocapsid (N) gene RNA. The SARS-CoV-2 *N* gene was PCR amplified from plasmids pUC57-SARS-CoV-2-N (GenScript) using forward primers with T7 promoter and reverse primers with poly(T)34 sequences. The *N* gene PCR product was gel purified and used as the template for *in vitro* transcription using the same mMESSAGE mMACHINE T7 Transcription Kit with 1 μg of DNA template, 1 μL of supplemental GTP in a 20 μL reaction volume. For SARS-CoV-2 IVT RNA electroporation, low passage BHK-21 cells were trypsinized and resuspended in cold PBS as 0.5×10^7^ cells/mL. A total of 20 μg of SARS-CoV-2 IVT RNA and 20 μg of *N* gene *in vitro* transcript were added to resuspended BHK-21 cells in a 2 mm gap cuvettes and electroporated with setting at 850 V, 25 μF, and infinite resistance for three times with about 5 seconds interval in between pulses. The electroporated cells were allowed to rest for 10 minutes at room temperature and were then co-cultured with Vero-hACE2-hTMPRSS2 cell at a 1:1 ratio in a T75 culture flask. Cell culture medium was changed to DMEM with 2% FBS the next day. Cytopathic effect (CPE) was monitored for five days, and cell culture supernatant were harvested for virus titration.

### Propagation of a clinical isolate of SARS-CoV-2 containing a deletion of the s2m

As part of ongoing SARS-CoV-2 variant surveillance, a random set of RT-PCR positive respiratory secretions from the Barnes Jewish Hospital Clinical microbiology laboratory were subjected to whole genome sequencing using the ARTIC primer amplicon strategy ^31^. This study was approved by the Washington University Human Research Protection Office (#202004259). From the sequences generated, one genome (2019-nCoV/WUSTL_000226/2020; Genbank # OM831956) had a 27-nucleotide deletion that removed 22 nucleotides from the 3’ end of the s2m element (**Fig S4**). The spike protein of this virus harbored L18F, D614G, and E780Q mutations, demonstrating that this virus belonged the original B.1 lineage of SARS-CoV-2. This virus was expanded twice on Vero-hACE2-hTMPRSS2 cells and the virus titer was determined by plaque assay. The P2 of 2019-nCoV/WUSTL_000226/2020 was sequenced by NGS to confirm the presence of the s2m deletion and rule out any tissue culture adaptations in the rest of the genome.

### SARS-CoV-2 growth curve and titration assays

Vero-hTMPRSS2 cells were grown to confluency. Cells were inoculated with a multiplicity of infection (MOI) of 0.001 of recombinant CoV-2-s2m-WT or mutant SARS-CoV-2 and culture supernatant was collected at 1, 24, 48, and 72 hpi and saved for viral quantification by plaque assay on Vero-hACE2-hTMPRSS2 cells in 24-well plates as described ^32^.

### SARS-CoV-2 Syrian hamster infection model

Animal studies were carried out in accordance with the recommendations in the Guide for the Care and Use of Laboratory Animals of the National Institutes of Health. The protocols were approved by the Institutional Animal Care and Use Committee at the Washington University School of Medicine (assurance number A3381–01). Five to six-week-old male hamsters were obtained from Charles River Laboratories and housed at Washington University. Next, the animals were challenged via intranasal route with 1,000 PFU of the recombinant CoV-2-s2m-WT or s2m mutant SARS-CoV-2 viruses under enhanced BSL-3 conditions^33^. Animal weights were measured daily for the duration of the experiment. Three and six days after the inoculation, the animals were sacrificed, and lung tissues and nasal washes were collected. The nasal wash was performed with 1.0 mL of PBS containing 1% BSA, clarified by centrifugation for 10 minutes at 2,000 x g and stored at −80°C. The left lung lobe was homogenized in 1.0 mL DMEM, clarified by centrifugation (1,000 x g for 5 minutes) and used for viral titer analysis by plaque assay and RT-qPCR using primers and probes targeting the *N* gene. For viral RNA quantification, RNA was extracted using RNA isolation kit (Omega Bio-tek). SARS-CoV-2 RNA levels were measured by one-step quantitative reverse transcriptase PCR (RT-qPCR) TaqMan assay as described previously using a SARS-CoV-2 nucleocapsid (N) specific primers/probe set from the Centers for Disease Control and Prevention (F primer: GACCCCAAAATCAGCGAAAT; R primer: TCTGGTTACTGCCAGTTGAATCTG; probe: 5’-FAM/ ACCCCGCATTACGTTTGGTGGACC/3’-ZEN/IBFQ)^34^. Viral RNA was expressed as (N) gene copy numbers per mg for lung tissue homogenates or mL for nasal swabs, based on a standard included in the assay, which was created via *in vitro* transcription of a synthetic DNA molecule containing the target region of the N gene.

### SARS-CoV-2 competetion assays in Syrian hamster

Ten golden Syrian hamsters were inoculated intranasally with a total of 1,000 PFU of a 1:1 mixture of CoV-2-s2m-WT and CoV-2-s2m^Δ2-42^ in 100 μL volume. Three days after inoculation, the hamsters were sacrificed, and the nasal washes, left lobes and lungs were harvested and homogenized. One hundred μL of the nasal wash or lung homogenates were added in 300 μL of TRK lysis buffer from E.Z.N.A Total RNA Kit (Omega Bio-tek) for RNA isolation. For quantitation of the virion-associated RNA levels in the inocula, we prepared five inocula separately and treated with RNase A (Thermo Scientific) to remove free viral genomic RNAs and subgenomic RNAs. These five RNase-treated inocula were added to TRK lysis buffer for RNA isolation and quantification as below to get the initial mutant to CoV-2-s2m-WT ratio. cDNA was synthesized from the extracted RNA with random hexamers using SuperScript IV First-Strand Synthesis Kit (Invitrogen) following the manufacturer’s protocol. PCR covering the virus S2M region was performed on cDNA samples for 40 cycles with primers HJ551-S2UTRF: 5’-CTCCAAACAATTGCAACAATC-3’ and HJ552-S2UTRR: 5’-GTCATTCTCCTAAGAAGCTATTAAAATC-3’ using the High Fidelity AccuPrime *Taq* DNA Polymerase (Invitrogen) following the manufacturer’s protocol. The presence of two different-size amplicons (389 bp for CoV-2-s2m-WT virus, 348 bp for CoV-2-s2m^Δ2-42^ virus) was verified by gel electrophoresis, and then PCR products were purified with QIAquick PCR Purification Kit (QIAGEN) and quantified with Qubit 4 Fluorometer (Invitrogen). Clean PCR products were diluted and subjected to 2100 Bioanalyzer (Agilent) analysis following the manufacturer’s protocol to get the molar quantities of the two different-size amplicons. Mutant to CoV-2-s2m-WT ratios were calculated based on quantitation readout, and the relative replicative fitness is defined and calculated by dividing the final mutant to CoV-2-s2m-WT ratio in hamster samples by the initial mutant to CoV-2-s2m-WT ratio in the inocula.

### Structural analysis

Vero-hTMPRSS2 cells were grown to confluency and the cells were inoculated with a MOI of 0.01 of recombinant CoV-2-s2m-WT or mutant CoV-2-s2m^Δ2-42^ virus. Supernatant was collected 24 hpi and the virus was purified from the supernatant using PEG preciptation method^35^. The purified virus pellet was resuspended in a buffer (0.05 M HEPES, pH 8, 0.1 M NaCl, 0.0001 M EDTA for SHAPE-MaP and 0.3 M HEPES, pH 8, 0.1 M NaCl for DMS-MaPseq).

#### SHAPE-MaP

For selective 2’-hydroxyl acylation analyzed by primer extension and mutational profiling (SHAPE-MaP), the resuspended viruses were divided into 3 reactions (modified sample, control sample, and denatured sample). For modified sample, 2-methylnicotinic acid imidazolide (NAI) from 1M stock (in DMSO) was added at a final concentration of 100 mM. For the control sample, the corresponding amount of DMSO was added. Samples were then incubated at 37° C for 15 min, followed by quenching of NAI through the addition of DTT at a final concentration of 0.5 M. For denaturing control, resuspended viruses were set aside without any treatment. TRK lysis buffer was added to above reactions to lyse the virus. Total RNA was extracted using Zymo RNA Clean and Concentrator-5 Kit (Zymo Research). The denatured control RNA sample was incubated at 95°C for 1 minutes and then treated with 100 mM NAI for 1 minutes at 95°C and the reaction was quenched with DTT as described previously. For denatured sample, RNA was again purified using Zymo RNA Clean and Concentrator-5 Kit (Zymo Research). Sequencing library preparation was performed according to the amplicon workflow as described previously^36^. The primers were designed tiling 3’ UTR across SARS-CoV-2 genome: 5’-GCAGACCACACAAGGC-3’ (forward) and 5’-CGTCATTCTCCTAAGAAGCTA-3’ (reverse). The RNA was reverse-transcribed using the specific primer with SuperScript II (Invitrogen) in MaP buffer (50 mM Tris-HCl (pH 8.0), 75 mM KCl, 6 mM MnCl_2_, 10 mM DTT and 0.5 mM deoxynucleoside triphosphate). Amplicons tiling the 3’ UTR SARS-CoV-2 genome were generated using Q5 hot start high-fidelity DNA polymerase (Cat. No. M0492S), 3’ UTR-specific forward and reverse PCR primers, and 8 μL of purified cDNA. The Nextera XT DNA Library Preparation Kit (Illumina) was used to prepare the sequencing libraries. Final PCR amplification products were size-selected using Agencourt AMPure XP Beads (Beckman Coulter). Libraries were quantified using a Qubit dsDNA HS Assay Kit (ThermoFisher, Cat. No. Q32851) to determine the concentration and quality was assessed with the Agilent High Sensitivity DNA kit (Agilent Technologies) on a Bioanalyzer 2100 System (Agilent Technologies) to determine average library member size and accurate concentration. The libraries were sequenced (2 × 150 base pairs (bp)) on a MiniSeq System (Illumina). Sequencing reads were aligned with the reference sequences and SHAPE-MaP reactivity profiles for each position was calculated using ‘Shapemapper-2.15’^37^ with default parameter. All SHAPE-MaP reactivities were normalized to an approximate 0–2 scale by dividing the SHAPE-MaP reactivity values by the mean reactivity of the 10% most highly reactive nucleotides after excluding outliers (defined as nucleotides with reactivity values that are >1.5 the interquartile range). High SHAPE-MaP reactivities above 0.7 indicate more flexible (that is, single-stranded) regions of RNA and low SHAPE-MaP reactivities below 0.3 indicate more structurally constrained (that is, base-paired) regions of RNA.

#### DMS-MaPseq

For dimethyl sulfate mutational profiling and sequencing (DMS-MaPseq), the resuspended viruses were divided into 2 reactions (modified and control sample). For modified sample, 2% v/v DMS was added, mixed thoroughly and incubated immediately at 37° C for 5 min before quenching with 100 μL 30% β-mercaptoethanol in PBS. The control sample was prepared similarly only without addition of DMS. TRK lysis buffer was added to above reactions to lyse the virus. Total RNA were extracted using Zymo RNA Clean and Concentrator-5 Kit (Zymo Research). For reverse transcription, the 11.5 μL RNA were supplemented with 4 μL 5× first strand buffer (ThermoFisher Scientific), 1 μL 10 μM reverse primer, 1 μL dNTP, 1 μL 0.1 M DTT, 1 μL RNaseOUT and 0.5 μL MarathonRT. The reverse-transcription reaction were incubated at 42° C for 3 hours. 1 μL RNase H was added to each reaction and incubated at 37°C for 20 min to degrade the RNA. cDNA was purified using QIAquick PCR Purification Kit (Cat. No. 28104). dsDNA were prepared as described above but with cocktail of 2 forward primers: 5’-GCAGACCACACAAGGC-3’ and 5’-ACGTTTTCGCTTTTCCG-3’. NEBNext® Ultra™ II DNALibrary Prep Kit for Illumina® (New England Biolabs, cat. E7645S) was used to prepare the sequencing libraries as per manufacturer instructions. The cleanup, quantification and sequencing of libraries was performed as described above. DMS-MaPseq reactivity profiles are calculated by aligning the sequencing reads to reference sequences using DREEM Webserver^38^. FastQC to assess the quality of fastq files, Trim Galore to remove adapter sequence and Bowtie2 to align the reads to sequnece are integrated into the DREEM. The DREEM map the reads and converts them into bitvectors based on the mutation rate (if mutation rate > 0.5%, converts to 1; otherwise, matches convert to 0). The bitvector files are then used to count mutations and normalized by sequencing depth in order to provide normalized DMS reactivity.

#### Experimentally-informed secondary structure modeling

To computationally predict the RNA secondary structure of the 3’ UTR, we have removed the primer-binding sequences from our analysis. The SHAPE-MaP and DMS-MaPseq reactivity profiles obtained through Shapemapper and the DREEM Webserver respectively, were used as constraints to predict the secondary structure using ShapeKnots^39^ with default settings. In order to calculate the correlation between the WT and mutant, the pearson correlation coefficient was calculated by comparing their nucleotide reactivity. SuperFold^40^ with experimental data restraints was used to predict consensus secondary structure prediction with base-pairing probabilities and Shannon entropy. The FASTA files for SHAPE-MaP and DMS-MaPseq can be found here: PRJNA936272

### Sequencing of reverse genetics rescued SARS-CoV-2 virus

For SARS-CoV-2 genome sequencing, 200 μL of supernatant containing virus was collected and added to 600 μL TRK lysis buffer plus beta-mecaptoethanol (BME) according to the E.Z.N.A. Total RNA Kit I (Omega Bio-tek). Total RNA was extracted according to the manufacturer’s protocol and eluted into RNase/DNase free H_2_O and used for library construction. Ribosomal RNA was removed from total RNA by Ribo-Zero depletion (Illumina). Indexed sequencing libraries were prepared using TruSeq RNA library preparation kit (Illumina), pooled and then sequenced using Illumina NextSeq system. The raw sequence data were analyzed using the LoFreq pipeline to call the mutations in the entire virus genome ^41^. In brief, Illumina sequencing fastq data was aligned by BWA with the SARS-CoV-2 reference genome sequence (NC_045512.2) after indexing to generate the aligned sam and bam files. Read group was added by Picard after the aligned bam file sorting and indexing with SAMtools, and then duplicates were removed by Picard MarkDuplicates. Local realignment was achieved by gatk3 and then variant was called by LoFreq to generate the mutant report file.

### Multiple sequence alignment and phylogenetic tree construction

Representative s2m sequences were aligned in MegaX using Muscle ^42^. Sequences were visualized using Jalview 2 ^43^. Representative sequences included: SARS-CoV-1 Urbani (AY278741.1), SARS-CoV-2 (NC_045512.2), Bat coronavirus RaTG13 (MN996532.2), Bat coronavirus BANAL-52 (MZ937000.1), Bat SARS coronavirus Rp3 (DQ071615.1), Bat SARS-like coronavirus YNLF-34C (KP886809.1), SARS coronavirus Civet PC4-136 (AY613949.1), Pangolin coronavirus PCoV GX-P3B (MT072865.1), Avian infectious bronchitis virus (NC_001451.1), Thrush coronavirus HKU12-600 (NC_011549.1). Representative SARS-CoV-2 complete genome sequences for each pango lineage were downloaded from NCBI. Sequence alignment was done by MAFFT and the phylogenetic tree was constructed with FastTree tool.

### Analysis of SARS-CoV-2 s2m sequences using the NCBI database

SARS-CoV-2 complete genome sequences deposited between January 2020 and December 2022 were downloaded from NCBI as a single FASTA file. The FASTA dataset was processed with a published SARS-CoV-2-freebayes pipeline to call all the variants along the genome ^44^. In brief, the single SARS-CoV-2 complete genome FASTA file was decomposed into individual FASTA genome sequences. Then, each FASTA genome was aligned individually against the SARS-CoV-2 reference sequence (NC_045512.2) using Minimap2 ^45^. Variant calling was performed on each BAM file using Freebayes variant caller to produce the VCF files. Variants in the s2m position 32 and deletion of the s2m Δ8-33 for all the VCF files were extracted and reported.

### Statistical analysis

Data was graphed using Prism 9.3.1 (GraphPad). Parametric and non-parametric comparisons were made where appropriate. Post hoc testing of Kruskal Wallis comparisons was completed with correction of multiple comparisons by Dunn’s multiple test correction. For multi-step growth curves, mutants were compared to CoV-2-s2m-WT by logarithmically transforming the data and comparing using a two-way ANOVA in Prism after excluding the one hour timepoint. Hamster weight data analyzed by mixed-effect model with Geisser-Greenhouse correction in Prism. Adjusted P values ≤ 0.05 were considered significant.

## AUTHOR CONTRIBUTIONS

H.J., A.J., C.F., T.Y., T.L.B., T.L.D. and H.H. performed experiments. A.J. and T.L.B performed the RNA probing studies and A.J. determined the RNA secondary structure of the 3’ UTR of SARS-CoV-2. B.P. obtained clinical isolates for local sequence analysis. S.T., A.J., J.A.B., B.F., and S.A.H. developed the sequence pipeline used for whole genome sequencing and performed sequencing of clinical and recombinant viruses. K.S. performed viral load analysis by RT-qPCR. T.L.B. and T.L.D. performed viral load analysis by plaque assay. H.J. and H.C. performed genetic analysis of the SARS-CoV-2 genomes. D.W. and A.C.M.B. provided supervision and acquired funding. H.J., A.J., A.B.J., D.W., and A.C.M.B. wrote the initial draft, with the other authors providing editorial comments.

## SUPPLEMENTARY INFORMATION

**Supplemental Figure 1:**
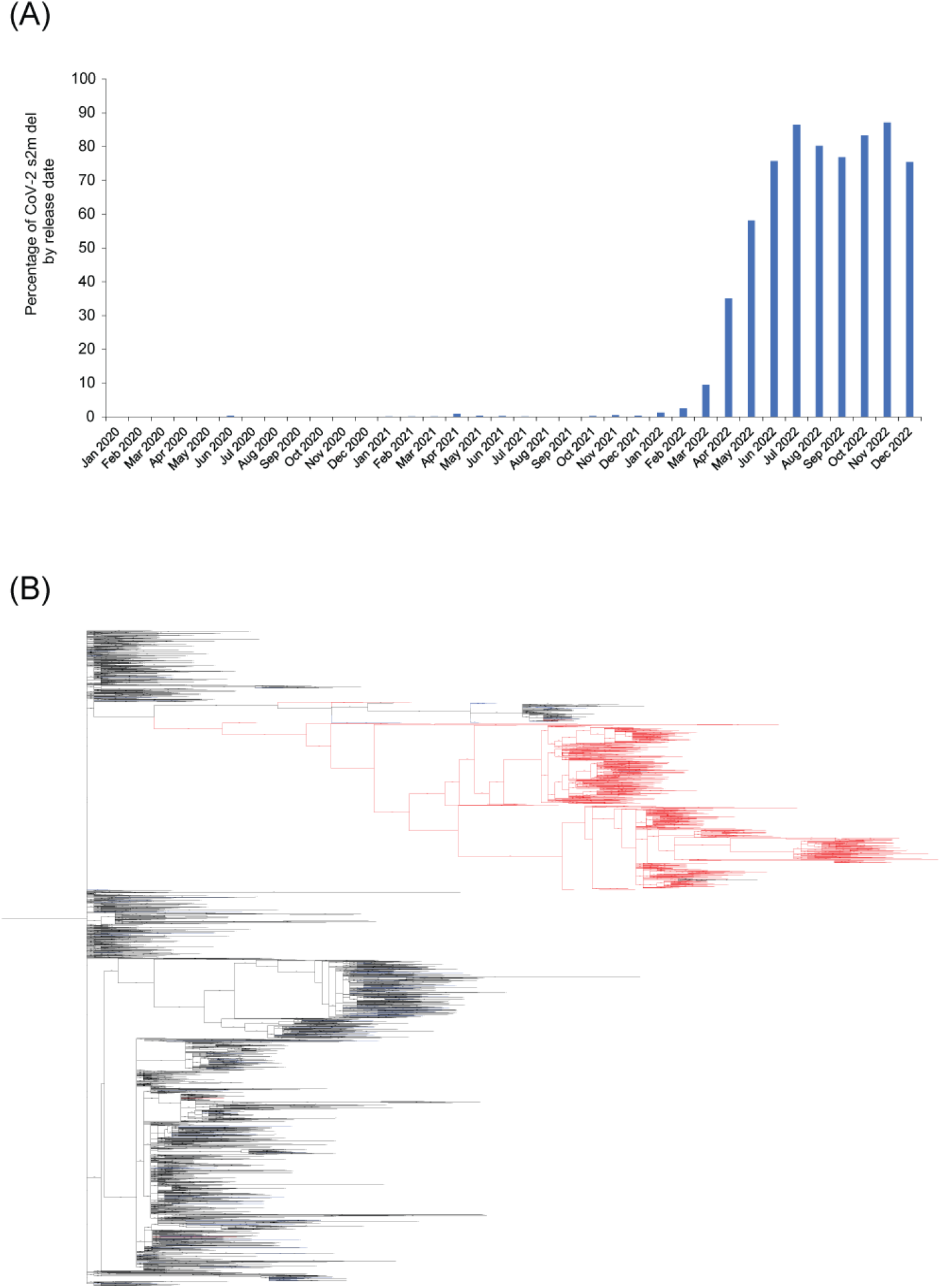
Identification of SARS-CoV-2 s2m deletions. (**A**) SARS-CoV-2 s2m deletion containing sequences found by release date. (**B**) A phylogenetic tree of SARS-CoV-2 by all Pango Lineages, SARS-CoV-2-s2m ^Δ8-33^ deletion containing lineages were marked as red. All other deletions in the s2m region are marked by blue.

**Supplemental Figure 2:**
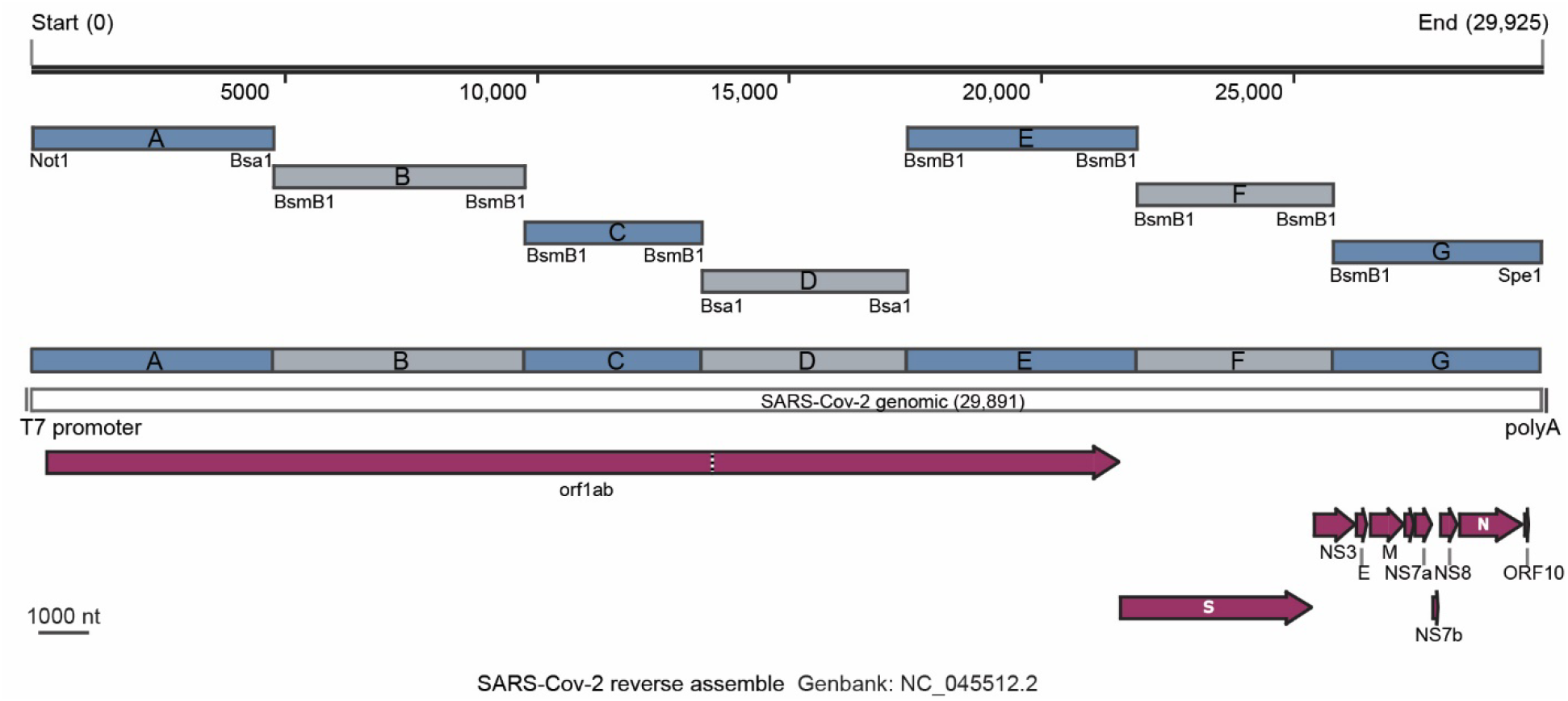
Schematic of the SARS-CoV-2 reverse genetics system used in this study. Schematic of the SARS-CoV-2 genome and fragment construction for the cDNA genome assembly. T7 promoter, polyA sequence, SARS-CoV-2 encoded ORFs were indicated on the genome map in purple. Restriction sites were shown in its position along the genome on each fragment.

**Supplemental Figure 3:**
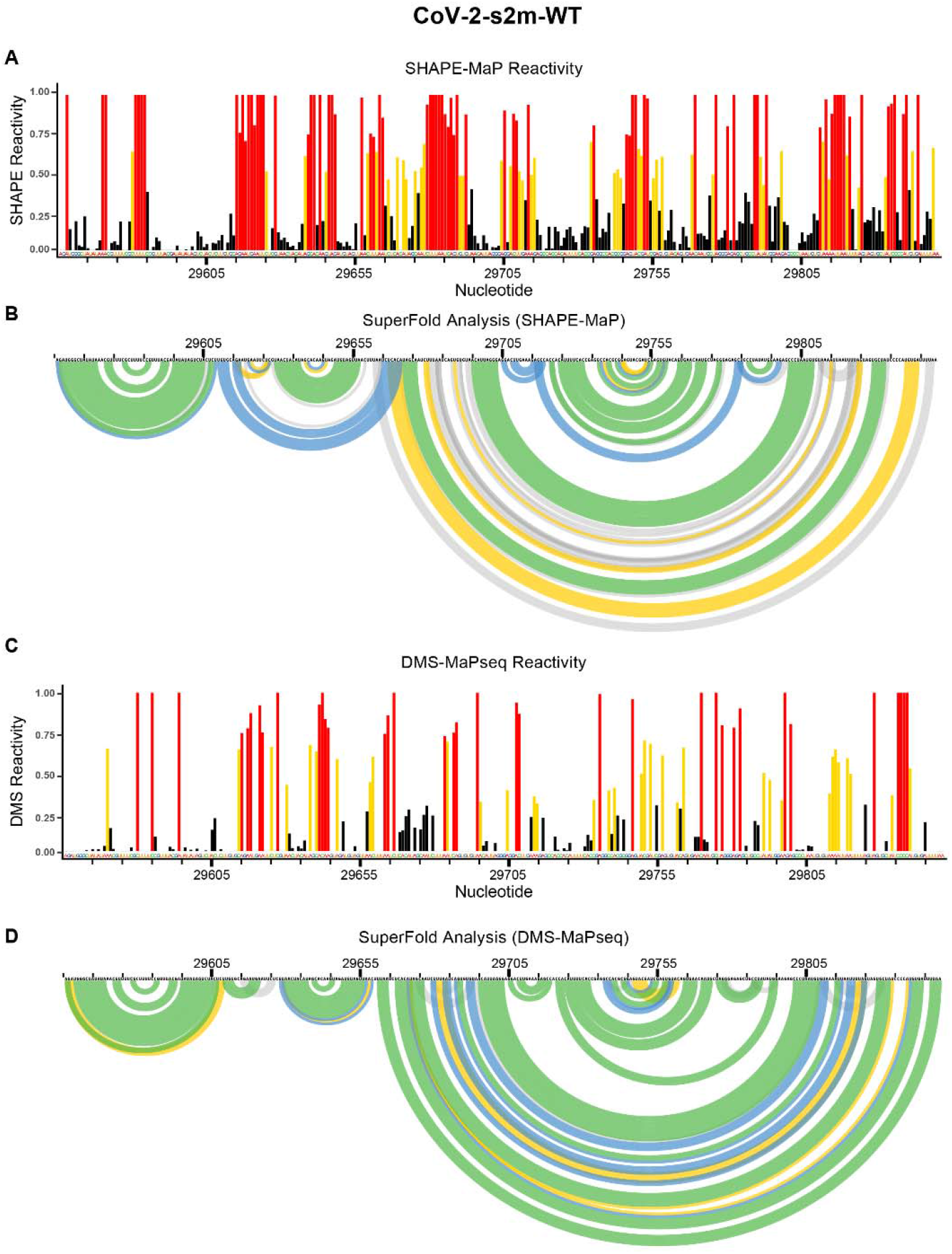
SHAPE-MaP and DMS-MaPseq reactivity profile and Superfold analysis of the 3’ UTR of wild type SARS-CoV-2. Following the probing of viral genome in purified SARS-CoV-2 virions, the SHAPE-MaP (A) and DMS-MaPseq (C) reactivity profiles were determined as described in the Materials and Methods section. SuperFold was used to predict consensus secondary structure prediction with base-pairing probabilities using the SHAPE-MaP (B) and DMS-MaPseq (D) reactivity profiles.

**Supplemental Figure 4:**
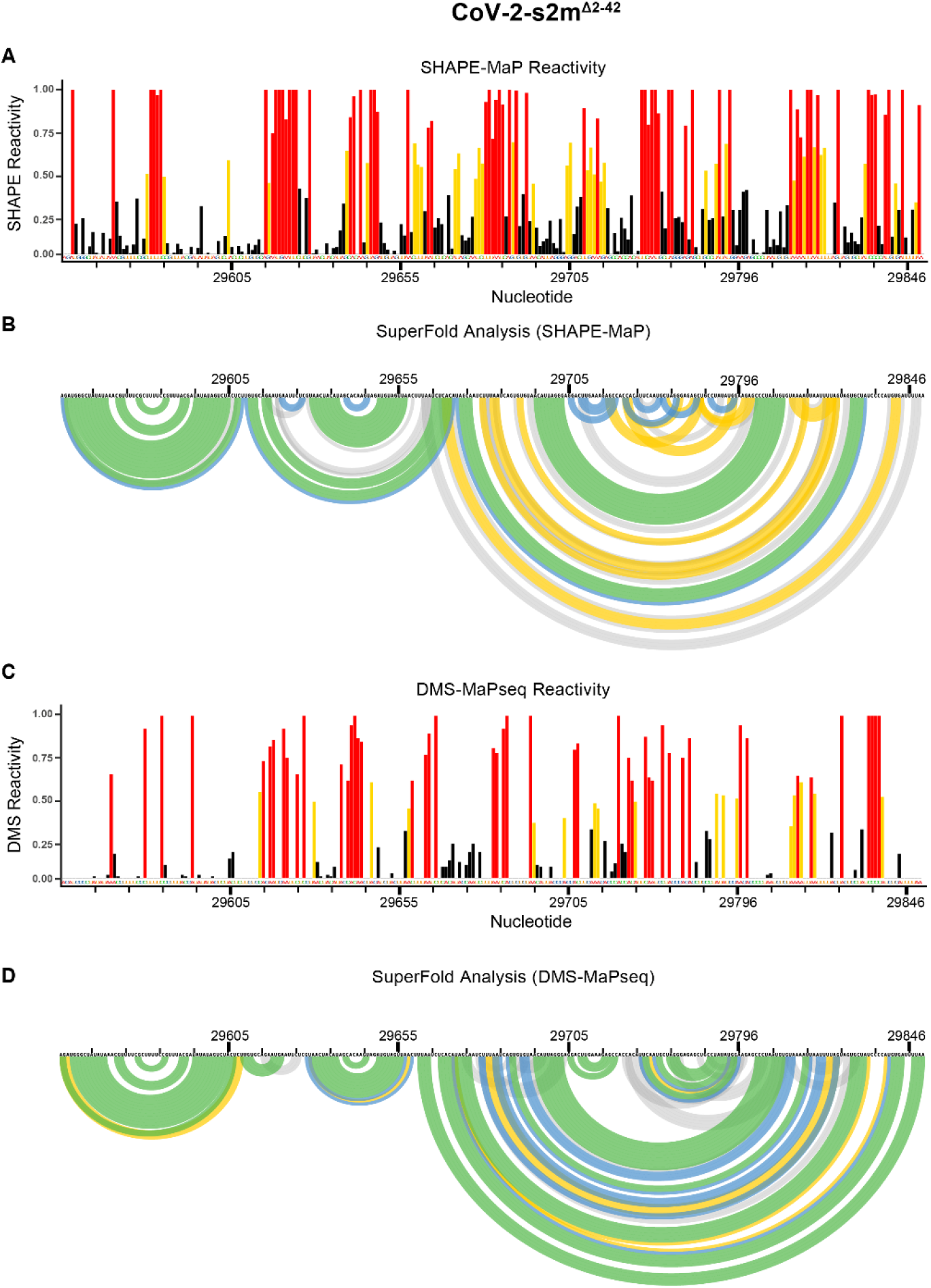
SHAPE-MaP and DMS-MaPseq reactivity profile and Superfold analysis of the 3’ UTR of SARS-CoV-2-s2m^Δ2-42^. Following the probing of viral genome in purified SARS-CoV-2 virions, the SHAPE-MaP (A) and DMS-MaPseq (C) reactivity profiles were determined as described in the Materials and Methods section. SuperFold was used to predict consensus secondary structure prediction with base-pairing probabilities using the SHAPE-MaP (B) and DMS-MaPseq (D) reactivity profiles. Nucleotides 29729 to 29768 were deleted from the CoV-2-s2m^Δ2-42^ genome.

